# Two sides to every coin: reciprocal introgression line populations in *Caenorhabditis elegans*

**DOI:** 10.1101/2022.08.29.505240

**Authors:** Mark G. Sterken, Lisa van Sluijs, Jelle W. van Creij, Daniel E. Cook, Joost A.G. Riksen, Katharina Jovic, Jasmijn Schouten, Maarten Steeghs, Yiru A. Wang, Jana J. Stastna, L. Basten Snoek, Simon C. Harvey, Jan E. Kammenga

## Abstract

Quantitative genetics seeks to understand the role of allelic variation in trait differences. Introgression lines (ILs) contain a single genetic locus introgressed into another genetic background, and are one of the most powerful quantitative trait locus (QTL) mapping designs. However, albeit useful for QTL discovery, this homogenous background confounds genetic interactions. Here, we created an IL population with N2 segments in a CB4856 background (IL_CB4856_), reciprocal to an N2 background with CB4856 introgressions population (IL_N2_). The IL_CB4856_ panel comprises a population of 145 strains with sequencing confirmed N2 introgressions in a CB4856 background. A core set of 87 strains covering the entire genome was selected. We present three experiments demonstrating the power of the reciprocal IL panels. First, we performed QTL mapping identifying new regions associated with lifespan. Second, the existence of opposite-effect loci regulating heat-stress survival is demonstrated. Third, by combining IL_N2_ and IL_CB4856_ strains, an interacting expression QTL was uncovered. In conclusion, the reciprocal IL panels are a unique and ready-to-use resource to identify, resolve, and refine complex trait architectures in *C. elegans*.

## Introduction

In the last decade, many advances have been made in quantitative genetics using *Caenorhabditis elegans*, putting it at the forefront of the study of natural genetic variation (EVANS *et al*. 2021). The most frequently used strains for exploring natural genetic variation in *C. elegans* are called N2 (Bristol) and CB4856 (Hawaii). These two strains differ in 176,543 single nucleotide variants 256,747 insertions-deletions (THOMPSON *et al*. 2015; KIM *et al*. 2019; LEE *et al*. 2021). These polymorphisms also translate to differences in phenotypic traits, many of which were identified using genetic crosses (EVANS *et al*. 2021). Many of these genes were identified using a combination of recombinant inbred lines (RILs) and introgression lines (ILs). These discoveries illustrate how ILs are excellent tools for dissection of complex traits. Importantly, ILs allow exploration of QTL effect sizes in both parental backgrounds. However, only a genome-wide IL panel in a single background exists for *C. elegans* at this time. Therefore we created a second genome-wide IL panel that is reciprocal to the existing IL panel.

Many quantitative genetics studies in *C. elegans* that focus on gene identification start with a two-step approach: first recombinant inbred lines (RILs) are used, followed by introgression lines (ILs) (ANDERSEN AND ROCKMAN 2022). RILs are lines that form a genetic mosaic of both parental genomes, whereas ILs consist of small segments of one parental strain introgressed into the genetic background of the other parental strain. In the RIL-IL approach, first a quantitative trait locus (QTL) is identified using RILs, followed by confirmation and fine mapping using ILs (for examples, see *e.g.*: (KAMMENGA *et al*. 2007; MCGRATH *et al*. 2009; BENDESKY *et al*. 2011; FREZAL *et al*. 2018; ANDERSEN AND ROCKMAN 2022) and for a more complete explanation see {Andersen, 2022 #3}). Although this paradigm is used in many QTL studies where such populations can be established, different approaches do exist such as chromosome-substitution populations or using whole-genome IL populations for trait mapping (KEURENTJES *et al*. 2007a; SCHMALENBACH *et al*. 2008; DOROSZUK *et al*. 2009; GLAUSER *et al*. 2011; FLETCHER *et al*. 2013). In studies using whole-genome IL populations, it has generally been noted that ILs are more sensitive for small-effect QTL compared to RILs (KEURENTJES *et al*. 2007a; GREEN *et al*. 2013; GLATER *et al*. 2014; BERNSTEIN AND ROCKMAN 2016; EVANS AND ANDERSEN 2020; EVANS *et al*. 2020). The reason ILs are more sensitive lies in the reduction of genetic diversity which enables direct comparison of isolated genomic regions against the parent background; genetic variation is isolated to the introgression. However, whole-genome ILs panels are generally less informative on the exact location of the QTL, because they contain fewer genetic cross-overs than RIL panels.

One of the persistent observations in quantitative genetics, is the occurrence of background-dependent QTL-effects. In other words, a locus can produce a different effect in an IL than in a RIL panel (for a review see {Kammenga, 2017 #276}). In pursuit of causal alleles/genes it is important to know potential background-effects, as they determine whether a QTL can be discerned. ILs are suitable for testing background effects. However, in a single- background introgression line population, the background-effects are by definition entangled. Therefore, IL populations with two backgrounds (containing introgressions from parent A in genetic background B and vice versa) would be of great value here. In combination reciprocal IL populations, loci-background interactions can be explored. Thus, adding a population with the reciprocal genetic background can further elucidate the genetic architecture of traits, as has already been demonstrated for specific QTL (*e.g.* (MCGRATH *et al*. 2009; GAERTNER *et al*. 2012; ANDERSEN *et al*. 2014)).

Here, we constructed a new IL population in *C. elegans* with N2 segments introgressed into a CB4856 genetic background (IL_CB4856_). This population complements a previously constructed IL_N2_ population and consists of 145 strains with sequencing-confirmed introgressions and a core set of 87 strains covering the entire genome. This makes *C. elegans* the first organism for which a set of reciprocal whole-genome IL panels is available. We present power simulations for QTL detection and show experimentally that this population can identify multiple QTL for lifespan (a notoriously complex trait). Furthermore, we demonstrate that the combination of the IL_CB4856_ and IL_N2_ panels can uncover genetic interactions with distant loci for heat-stress survival and the expression of the gene *clec-62*. Together, our research provides a toolset for uncovering and mapping the existence of complex genetic architectures in *C. elegans*.

## Material and methods

### Strains and maintenance

The starting strains for introgression line construction were recombinant inbred lines (RILs) with N2 and CB4856 parents, namely: WN001, WN007, WN025, WN068, WN071, and WN110 (LI *et al*. 2006). Strains were kept on NGM plates seeded with *Escherichia coli* OP50 and culturing temperatures used during the crosses were 12^°^C, 16^°^C, or 20^°^C, depending on the desired speed of population growth (BRENNER 1974). A subset of 15 of the IL_CB4856_ population – covering parts of chromosomes I and IV – has been published previously (GAO *et al*. 2018; STERKEN *et al*. 2021).

### Crossing scheme

To generate the IL_CB4856_, we divided crosses into two stages (**Supplementary table 1**). The first stage was used for most loci, where a RIL male was back-crossed to a CB4856 hermaphrodite to ensure the presence of CB4856 mitochondria in the F1. This step was followed by a second stage cross with CB4856 males to enable the integration of homozygous CB4856 genotypes at the *peel-1 zeel-1* incompatibility locus on chromosome I in the F2 (SEIDEL *et al*. 2011). The exception were strains with WN001 as a parent, where we wanted to obtain coverage of the *peel-1 zeel-1* locus. For WN001 crosses, the second cross was initially omitted. Subsequently, selected genotyping was conducted in the F3 (4-9 markers; **Supplementary table 2**), screening for strains with most CB4856 loci and absence of the N2 genotype at the *peel-1 zeel-1* locus.

Next, selected strains were inbred and selected further to obtain as many homozygous CB4856 loci as possible (for 5-12 generations). If a desired strain contained more than one N2 locus, further back-crosses with CB4856 males were conducted until only one detectable N2 locus remained in an otherwise CB4856 genotypic background. Finally, the strains were inbred by transferring single hermaphrodites to new NGM plates for at least 10 generations. In this way, 154 ILs were created initially. These ILs have been cryopreserved along with the parental strains. Ultimately, after genotyping by low-coverage whole-genome sequencing we could verify the introgressions in 145 IL_CB4856_ strains and selected a core set of 87 unique introgression lines covering the entire genome (**Supplementary table 3**).

### Genotyping by fragment length polymorphisms

Initial genotyping (during the crossing of the strains) was PCR-based using primer pairs that detect insertions-deletion variants between the CB4856 and N2 strains (THOMPSON *et al*. 2015). In total, 41 primer pairs were optimized with a bias for covering loci with a high-recombination frequency (**Supplementary table 2**) (ROCKMAN AND KRUGLYAK 2009). The selection criteria for generating the primer pairs were: (i) the deletion occurred in the CB4856 strain, (ii) the deletion is larger than 25 bp and shorter than 150 bp, and (iii) it is not located in a repetitive region. All primers have been developed with Primer3 (primer3-win-bin-2.3.6) on the 1000 bp up- and downstream of the deletion (UNTERGASSER *et al*. 2012). Primer3 was used with standard settings, selecting three primers in the size ranges of: 100-150 bp, 200-250 bp, 300-350 bp, 400-450 bp, 500-550 bp, 600-650 bp, 700-750 bp, and 800-850 bp. The annealing temperature was selected between 58^°^C and 60^°^C. The specificities of the primers were tested using BLAST (ncbi-blast 2.2.28 win64) against WS230 (settings: blastn – word_size 7 –reward 1 –penalty -3) (ALTSCHUL *et al*. 1990). Only primers with fewer than five hits were considered for further selection. Final selection of the primers was based on the presence of a visible amplicon of the expected size after PCR of the N2 and CB4856 strains (**Supplementary figure 1**) and detection of heterozygous strains.

For genotyping during crossing, DNA was isolated from single adults that had generated offspring. Nematodes were lysed at 65^°^C for 30 minutes using a custom lysis buffer (VERVOORT *et al*. 2012), followed by 5 minutes at 99^°^C. Genotyping PCRs were performed with GoTaq using the manufacturers recommendations (Promega, Catalogue No. M3008; Madison, USA). The annealing temperature was 58^°^C (30 seconds), with an extension time of 1 minute for 40 cycles. All samples were run on 1.5% agarose gels stained with ethidium bromide. Ultimately, a low-resolution genetic map was constructed for 154 ILs based on the insertion-deletion markers.

### Sequencing: DNA isolation, library construction, and sequencing

DNA was isolated from all the 154 initially generated IL_CB4856_. Furthermore, DNA was isolated from the six parental N2xCB4856 RILs: WN001, WN007, WN025, WN068, WN071, and WN110. We also sequenced an additional 29 IL_N2_ that were generated previously (DOROSZUK *et al*. 2009): WN203, WN204, WN206, WN208, WN210, WN213, WN214, WN218, WN222, WN224, WN231, WN233, WN236, WN237, WN238, WN240, WN247, WN249, WN253, WN256, WN259, WN261, WN262, WN265, WN269, WN272, WN282, WN283, and WN285. Furthermore, we sequenced the two parental strains N2 and CB4856 as reference (**Supplementary table 3**).

The DNA isolation and library construction have been reported on previously (GAO *et al*. 2018). Shortly, genomic DNA was obtained from populations grown on 9 cm NGM plates. The Qiagen DNeasy Blood & Tissue Kit (Catalogue No. 69506; Hilden, Germany) was used for DNA isolation. DNA was quantified by Qubit. For each sample, a total of 0.75 ng of DNA was taken as input for the library construction. The libraries were sequenced on an Illumina HiSeq using a 300 cycle kit. Data have been deposited under accession number SRP154243 in the NCBI Sequence Read Archive (SRA; https://www.ncbi.nlm.nih.gov/sra).

### Genotype calling using hidden Markov model and construction of the genetic map

The low-coverage sequencing data were used for variant calling using a hidden Markov model, as described previously (COOK *et al*. 2016; COOK AND ANDERSEN 2017). Subsequently, the hidden Markov model genotype calls were filtered for low-coverage areas (<100 supporting calls) and for introgressions less than 1*10^5^ base pairs, which are unlikely to occur often given the crossing scheme, and more likely to be incorrect calls. From the 154 initially created ILs, we could confirm the presence of an introgression in 145 ILs and a core set of 87 ILs was selected. The core set covered as many loci as possible with introgressions with as few multi-introgression ILs as possible. Ultimately, the genetic map of the new population and the additionally sequenced strains were integrated with previously constructed and sequenced RILs (LI *et al*. 2006; THOMPSON *et al*. 2015) IL_N2_ strains (DOROSZUK *et al*. 2009; THOMPSON *et al*. 2015) from the Laboratory of Nematology, Wageningen University. The map was constructed from the identified points of recombination within these populations. The markers describing the populations are adjacent to either side of each point of recombination. This led to a genetic map with 1152 markers covering the recombination events in the whole set of strains (**Supplementary table 4**).

### Lifespan experiment

We measured lifespan of the 87 IL_CB4856_ strains in the core set. Each experiment was started by transferring 10 L4 animals to three independent plates per genotype (30 animals per strain in total). The two parental strains were tested on nine plates (90 animals in total). Plates were screened daily until reproduction ended. Animals that showed egg laying deficiencies (also known as ‘bagging’) and those that were lost from the assay (i.e. that climbed to the sides of the plate and desiccated) were censored from the data. For the IL_CB4856_ strains WN322, WN342, and WN351, all three replicates did not yield enough observations. For QTL mapping, we summarized the data to the mean, median, minimum, maximum, and variance in lifespan. These summarized traits were used in further analyses (**Supplementary table 5**).

### Heat-shock survival experiment

We analysed previously published data to identify heat-stress survival QTL in *C. elegans* (JOVIC *et al*. 2019). The goal of the analysis was to test whether loci on chromosome IV could be associated with heat-stress survival. The data from Jovic *et al*. consist of survival measurements in 33 CB4856 x N2 RILs and 71 IL_N2_ strains (**Supplementary table 7**). For each strain, 32 animals were measured together on one plate on average. These animals were heat-shocked for four hours at 35^°^C, 48 hours after age-synchronization of the population by bleaching (see (JOVIC *et al*. 2019)). We took the data for survival at 72, 96, and 216 hours for mapping QTL.

As the previously published dataset indicated chromosome IV might be involved in heat-stress survival, we designed a replicated experiment. We tested 17 IL_N2_, 20 IL_CB4856_, and the N2 and CB4856 parental strains for heat-shock survival (**Supplementary table 8**). Each experiment was started by transferring a food-deprived population to a new 9 cm NGM dish, where the population was allowed to develop for ∼60h at 20^°^C. After that period, the population consisted of egg-laying adults, from which eggs were isolated by bleaching for developmental synchronization (day 0) (EMMONS *et al*. 1979). Isolated eggs were grown for 48 h at 20^°^C. At this time, 20-40 nematodes were transferred to 6 cm NGM plates containing 100 µl FUdR, which inhibits reproduction (HOSONO 1978). Two plates per strains were generated, one as a control (remaining on 20^°^C) and one receiving a four hour 35^°^C heat- shock immediately after transfer (RODRIGUEZ *et al*. 2012). At 72h, 96h and day 216h of the experiment, the number of surviving and dead nematodes were counted. After gathering the data, we only included samples with greater than 10 animals found at the 72 hour observation (upon heat-shock, animals tend to crawl to the top of the plates where they desiccate). Each strain was assessed at least three times per treatment, parental lines were tested more frequently (n = 10 for N2 and n = 12 for CB4856).

### Experiment to measure gene expression of clec-62

We used the IL_CB4856_ population to test the QTL for the expression of *clec-62*, for which multiple QTL were uncovered in a previous study in the IL_N2_ panel (STERKEN *et al*. 2020). Expression of *clec-62* was measured in a similar setup as in two previous eQTL experiments (SNOEK *et al*. 2017; STERKEN *et al*. 2020). In short, we collected 48-hour old L4 nematodes of N2 (6 replicates), CB4856 (6 replicates) and a single replicate for 46 IL_CB4856_ strains, 50 IL_N2_ strains, and 52 N2xCB4856 RILs grown at 20^°^C (**Supplementary table 9)**. The strains were tested in three experimental batches and each batch contained a randomized selection of strains and both parental lines in duplicate.

The RNA was isolated from these samples using a Maxwell16 LEV Plant RNA kit using the recommended protocol with one modification, namely adding 20 µL proteinase K to the lysis step after which the samples were incubated for 10 minutes at 65°C whilst shaking at 900 rpm (STERKEN *et al*. 2014). Some samples were removed from the analysis afterwards because of low RNA concentrations and/or RNA degradation (confirmed by gel electrophoresis). Subsequently, cDNA was constructed using a Promega GoScript reverse transcriptase kit and *clec-62* expression was measured using RT-qPCR. For the RT-qPCR, two primer pairs were designed for amplification of the two isoforms (A and B) of *clec-62:* P_MS_CLEC62_A_F (5’ CGACACTTCATTCCCCGAGC 3’), P_MS_CLEC62_A_R (5’TTAAGCTGGAACGGCACCAAC 3’), P_MS_CLEC62_B_F (5’CGCGTTGGTGCCGCTTAAC 3’) and P_MS_CLEC62_B_R (5’GATTGCTGATTGAGGACGGCG 3’). Gene expression of *clec-62* was normalized against the reference genes *rpl-6* and *Y37E3.8* (STERKEN *et al*. 2014). After the experiment and quality control of the qPCR data (sufficient amplification of the reference genes {Sterken, 2014 #428}), 5 N2 replicates, 6 CB4856 replicates, 41 IL_CB4856_, 42 IL_N2_, and 47 RIL samples were analysed further.

### QTL mapping in the IL populations

We used three approaches for QTL mapping in the IL population(s): (i) to an individual IL, (ii) to a single genetic background using bin mapping (DOROSZUK *et al*. 2009), and (iii) to two backgrounds using bin mapping with an interaction term. The mapping to an individual IL uses a linear model where each IL is compared to the parent genetic background, correlating trait differences to the introgression region covered by the particular IL. To correlate trait values to the introgression, the linear model

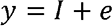

where *y* is the trait and *I* is the presence of the introgression was used. Each IL was tested with this model versus the genetic-background parent (*e.g*., for an IL_CB4856_ strain versus the CB4856 strain). The obtained p-values were corrected for multiple testing by using the p.adjust function in R with the Benjamini-Hochberg method (BENJAMINI AND HOCHBERG 1995).

The single-background bin mapping assumes that a single QTL exists per overlapping set of introgressions (DOROSZUK *et al*. 2009). This method is useful when all ILs covering a particular chromosome are tested. Here, the ILs with an introgression at a specific marker were tested versus the genetic background strain. Similar to the individual introgression, a linear model

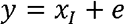

where *y* is the trait and *x_I_* is the introgression genotype at a given marker is used. For each marker, only the ILs with an introgression are compared to the genetic background parent (*e.g*., for an IL_CB4856_ strain versus the CB4856 strain). The significance threshold was calculated using a permutation approach with 1,000 permutations to determine whether a significance fell below a pre-set false-discovery rate (q = 0.05).

The two-background bin mapping also assumes that a single QTL exists per overlapping set of introgressions. However, because both parental backgrounds and the derived ILs were considered, the interaction between the two backgrounds can also be solved

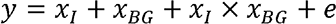

Where *y* is the trait and *x_I_* is the introgression genotype at a given marker, and *x_BG_* is the genetic background of the IL. Also here, for each marker, only the ILs with an introgression are compared to the genetic background parent (*e.g*., for an IL_CB4856_ strain versus the CB4856 strain). The significance threshold was calculated using a permutation approach with 1,000 permutations to determine whether a significance fell below a pre-set false-discovery rate (q = 0.05).

Using the outcome of the bin mapping models QTL peaks were called using a 1.5 LOD-drop and under the condition that two peaks should be at least 1*10^6^ base pairs apart.

### QTL mapping in the RIL population

Mapping in the RIL population was conducted as described previously (GAO *et al*. 2018; STERKEN *et al*. 2021). In short, we fitted the trait data to the linear model

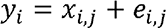

where *y* is the trait as measured in RIL *i*, *x* is the marker of RIL *i* at location *j*, and *e* is the residual variance. The 1152 marker set was used for mapping, and the significance was determined by a permutation approach using 1,000 permutations (q = 0.05).

### Power analysis in the IL populations

To determine the statistical power of bin mapping with the IL_CB4856_ and both IL panels, we used an *in silico* simulation approach. The scripts for the simulation can be found in the git archive (see below). QTL mapping was simulated for two scenarios: (i) using only the IL_CB4856_ panel (87 strains), or (ii) using the combined IL_CB4856_ and IL_N2_ panels. To test the power for QTL detection, simulated data was used as input for the analyses. First, we simulated scenarios of a single QTL, varying the amount of variance explained (0.2, 0.25, …, 0.8), this was simulated by adding normally distributed ‘noise’ to the dataset. We simulated these QTL per marker in the map (i.e. the QTL was directly located on the marker). We also simulated various levels of replication in the RILs and ILs (2, 3, …, 15), thus multiple measurements of the same trait that went into the mapping algorithms. QTL were mapped using the methods described in the previous two sections. The outcomes were compared to what was expected based on the simulation (i.e. where the QTL should be found versus where the QTL was actually mapped).

### Heritability analysis

The broad-sense heritability (*H^2^*) was calculated as in (BREM AND KRUGLYAK 2005; KEURENTJES *et al*. 2007b; STERKEN *et al*. 2020), where

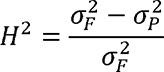

here *H^2^* was the broad-sense heritability, σ^2^ is the variance of either the population *F* (RIL, IL_N2_, or IL_CB4856_) or the parental strains *P* (N2 and CB4856). Where the variance of the parental populations was the pooled variance and used as an estimate of the measurement error. The significance threshold was determined using a permutation approach with 1,000 permutations per trait. A significance of q = 0.01 was taken to compensate for the upper-bound estimation this approach gives.

The narrow-sense heritability was calculated using a REML approach as provided by the “heritability” package (SPEED *et al*. 2012; KRUIJER AND KRUIJER 2014; KRUIJER *et al*. 2015). The significance threshold was determined by 1,000 permutations per trait.

### Software, scripts, and data

Data were analysed using R (version 3.4.2, windows x64) in RStudio (version 1.1.383) with custom written scripts (TEAM 2013; TEAM 2015). The tidyverse packages (version 1.2.1) were used for organizing and plotting data (WICKHAM 2019). All scrips and data are available at https://git.wur.nl/published_papers/sterken_2022_cb4856-ils. In particular, the scripts include a set of functions for QTL mapping and power simulations in the IL populations.

## Results

### Genetic characteristics of the IL_CB4856_ population

A genome-wide population of introgression lines containing an N2 segment in a CB4856 background was constructed (IL_CB4856_). This set was created by backcrossing a set of six recombinant inbred lines (LI *et al*. 2006) with the CB4856 strain (**Supplementary table 1; Figure 1A**). During the crosses, the genotypes were monitored using 41 amplification fragment length polymorphism (FLP) markers (**Supplementary table 2; Supplementary figure 1**) and afterwards all 154 selected IL strains were whole-genome sequenced (**Supplementary table 3**). A total of 145 strains with sequence-confirmed N2 introgressions into the CB4856 strain was obtained, of which 99 contained a single N2 introgressed region in an otherwise CB4856 genetic background, 37 ILs contained two N2 introgressed regions, and nine strains contained multiple N2 introgressed regions (**Supplementary table 3**). From this set, a core set of 87 strains was assembled, covering the entire *C. elegans* genome with N2 segments in an otherwise CB4856 genetic background (**Figure 1B**). The median introgression size of this population was 3.13 Mb, with the smallest region spanning less than 0.02 Mb and the largest region spanning 16.09 Mb. Each chromosome was covered by regions found in a minimum of seven ILs (chromosome III) and a maximum of 20 ILs (chromosome IV). In conclusion, this set of strains can be used as resource for QTL exploration in *C. elegans*.

**Figure 1:**
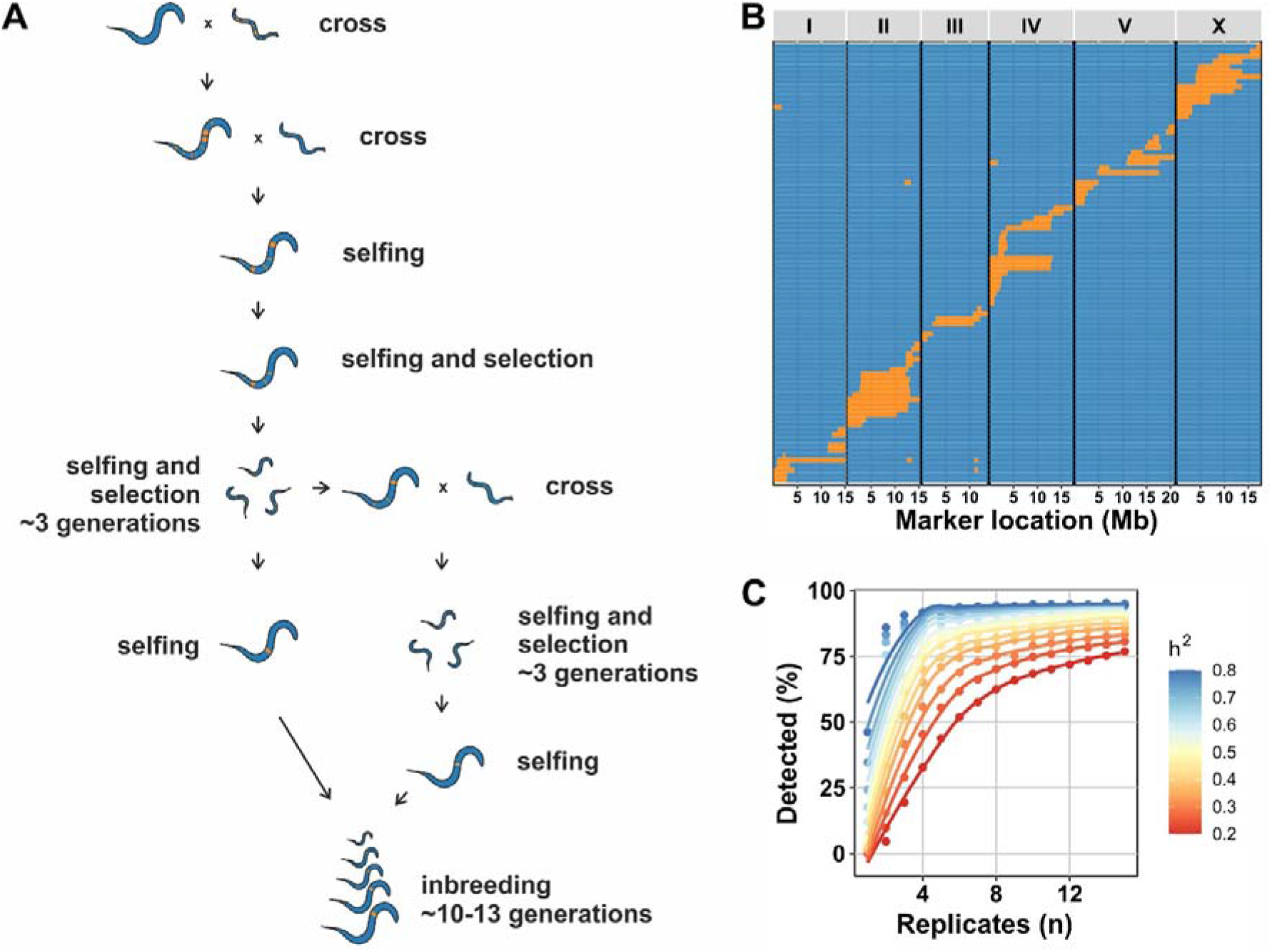
A novel N2>CB4856 introgression line panel and its mapping properties. (A) Schematic overview of the crossing scheme to create IL_CB4856_ strains. N2xCB4856 RIL strains were (back)crossed with CB4856 over several generations to obtain IL_CB4856_ strains with single introgressions. (B) The panel consisting of 87 strains covering the entire genome with N2 introgressions in a CB4856 genetic background. On the x-axis, the physical location is shown in million bases (Mb) split-out per chromosome. On the y-axis, the strains are shown (no label, each row represents a single strain). The blue colours indicate the CB4856 genotype, and the orange colours indicate the N2 genotype. (C) *In silico* power analysis of the detection of background interactions when combining the IL_CB4856_ with the previously constructed IL_N2_ panel. On the x-axis, the number of replicates per IL is plotted against the percentage of the simulated QTL detected on the y-axis. The colours indicate the amount of variance explained (*h^2^*) per simulated interaction QTL.

To facilitate integrated analyses using IL populations, we created and tested the power of a combined genetic map of the new population with our previously constructed N2- background IL population (IL_N2_) (DOROSZUK *et al*. 2009). To complete the IL_N2_ genetic map, we sequenced 29 of the IL_N2_ strains, to supplement the 57 IL_N2_ strains that were sequenced previously (THOMPSON *et al*. 2015). By integrating the IL_CB486_, IL_N2_ with the genetic map of the Wageningen N2xCB4856 RIL population (LI *et al*. 2006; THOMPSON *et al*. 2015), we constructed a map with 1152 informative markers, spanning 389 strains (**Supplementary table 4**). We used this map for power analyses of the core sets of the two IL populations. With the establishment of an IL_CB4856_ population, we could simulate its ability to detect QTL when combined with the IL_N2_ population (**Supplementary figure 2; Figure 1C**). Using simulations, we found that three replicates are sufficient to detect 50% of the interacting QTL explaining at least 35% of variance. Therefore, this analytical design can effectively uncover local genetic interactions.

### QTL mapping in the IL_CB4856_ panel uncovered nine lifespan QTL

To investigate QTL detection using real data in the newly generated IL_CB4856_ strains, we performed a QTL mapping experiment for lifespan (**Supplementary table 5**). We picked this trait because it was tested previously in the IL_N2_ population and would allow us to compare the QTL architectures (DOROSZUK *et al*. 2009). In the previous panel, six QTL for mean lifespan were detected. When we measured lifespan, we found that the N2 strain lived longer than the CB4856 strain on average, which was in line with previous results (**Figure 2A**) (DOROSZUK *et al*. 2009). After bin mapping, we detected 42 QTL for the various summary statistics mapped, including nine QTL related to mean lifespan (**Supplementary figure 3; Supplementary table 6**; **Figure 2B**). Of these nine mean-lifespan QTL, three (1, 2, and 7) were also detected in the IL_N2_ population {Doroszuk, 2009 #12}. For example, the effect direction of QTL 7 (shorter mean lifespan related to CB4856) was recapitulated in the IL_CB4856_ population (**Figure 2C**), indicating that we uncovered an additive QTL on the right side of chromosome IV.

**Figure 2:**
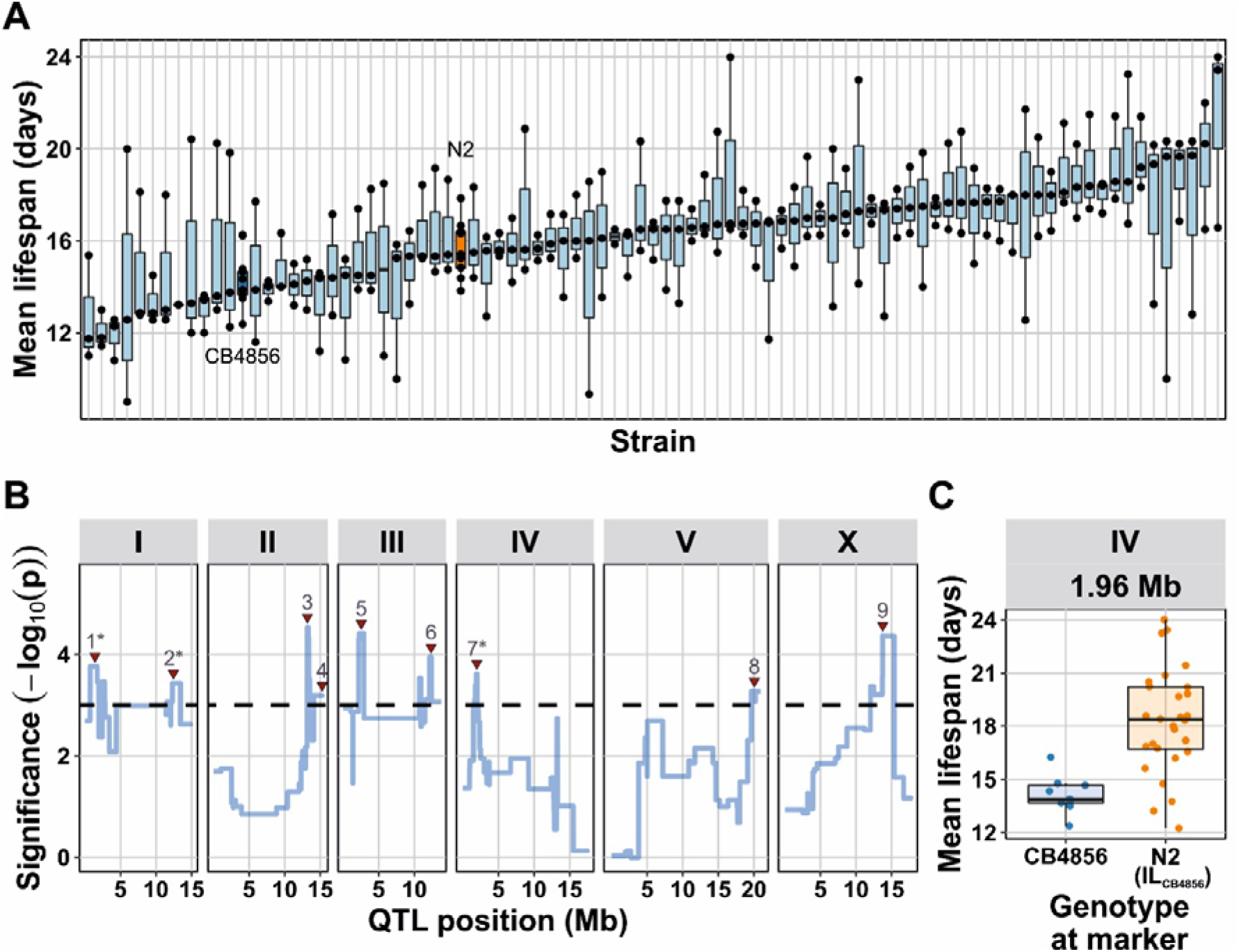
Lifespan analysis in the IL_CB4856_ panel. (**A**) Overview of the mean lifespan over all 87 IL_CB4856_ strains and the N2 and CB4856 parental lines. Each dot represents a plate (n = 10 animals per plate), the box plots are used as visual aids. The colours indicate the genetic background (orange: N2, blue CB4856). Most of the ILs had longer lifespan than the two parental lines. (**B**) The QTL profile for the mean lifespan mapping. On the x-axis, the physical position in million bases (Mb) is shown. On the y-axis, the statistical significance of the association. The dashed horizontal line indicates the FDR = 0.05 threshold (based on 1,000 permutations). The red triangles indicate the peaks that were called, the asterisk indicates that this peak was mapped previously in the IL_N2_ panel (**C**) The trait values under the chromosome IV peak (7), where we found that the ILs with an introgression on the site (N2 genotype) had an increased lifespan compared to the CB4856 parental strain.

### QTL fine mapping using the IL_CB4856_ panel identified 18 heat-stress survival QTL on chromosome IV

The most common usage of ILs is in fine mapping or experimental validation of previously identified QTL. Typically, once a QTL is detected in a RIL panel, a selected set of ILs can be used to validate a QTL detected in a mapping experiment (for examples, see *e.g.*: (Kammenga *et al*. 2007; McGrath *et al*. 2009; Bendesky *et al*. 2011; Frezal *et al*. 2018; ANDERSEN AND ROCKMAN 2022)). We used a set of ILs that could validate the detected QTL on chromosome IV in the heat-stress response. Chromosome IV was previously implicated in fitness (reproduction) after heat shock, and a heat-shock expression QTL hotspot was found on chromosome IV (RODRIGUEZ *et al*. 2012; SNOEK *et al*. 2017; STERKEN *et al*. 2020). To further verify the QTL, we performed a QTL analysis on previously published data from a heat-shock survival experiment in the IL_N2_ panel (**Supplementary table 7**) (JOVIC *et al*. 2019). In this experiment, we found only one chromosome IV QTL associated with heat-stress survival (**Supplementary figure 4; Supplementary table 6**). We set out to investigate (i) whether this QTL could be replicated and (ii) whether it was additive or implicated in a genetic background interaction.

We performed a heat-stress survival experiment on ILs that together covered chromosome IV: 17 IL_N2_ strains and 20 IL_CB4856_ strains. We measured survival of four hour exposure to 35°C at 24, 48, and 168 hours after the start of the exposure (**Supplementary table 8**; **Figure 3A**). We observed that the ILs typically showed a decreased survival when compared with the parental lines (**Figure 3B**). When we used these data for bin mapping, 18 heat-stress survival QTL were uncovered on chromosome IV (**Supplementary table 6; Figure 3C**). One locus, a QTL around 12.6 Mb was associated with a decrease in survival in both N2 and CB4856 ILs, this was also the QTL we mapped in the previously published data (**Supplementary figure 4**). Mapping in both IL populations indicated the existence of opposite-effect QTL in that region (**Figure 3D** and **E**). Therefore, we concluded that the use of strains from both IL panels is useful to uncover background-effects.

**Figure 3:**
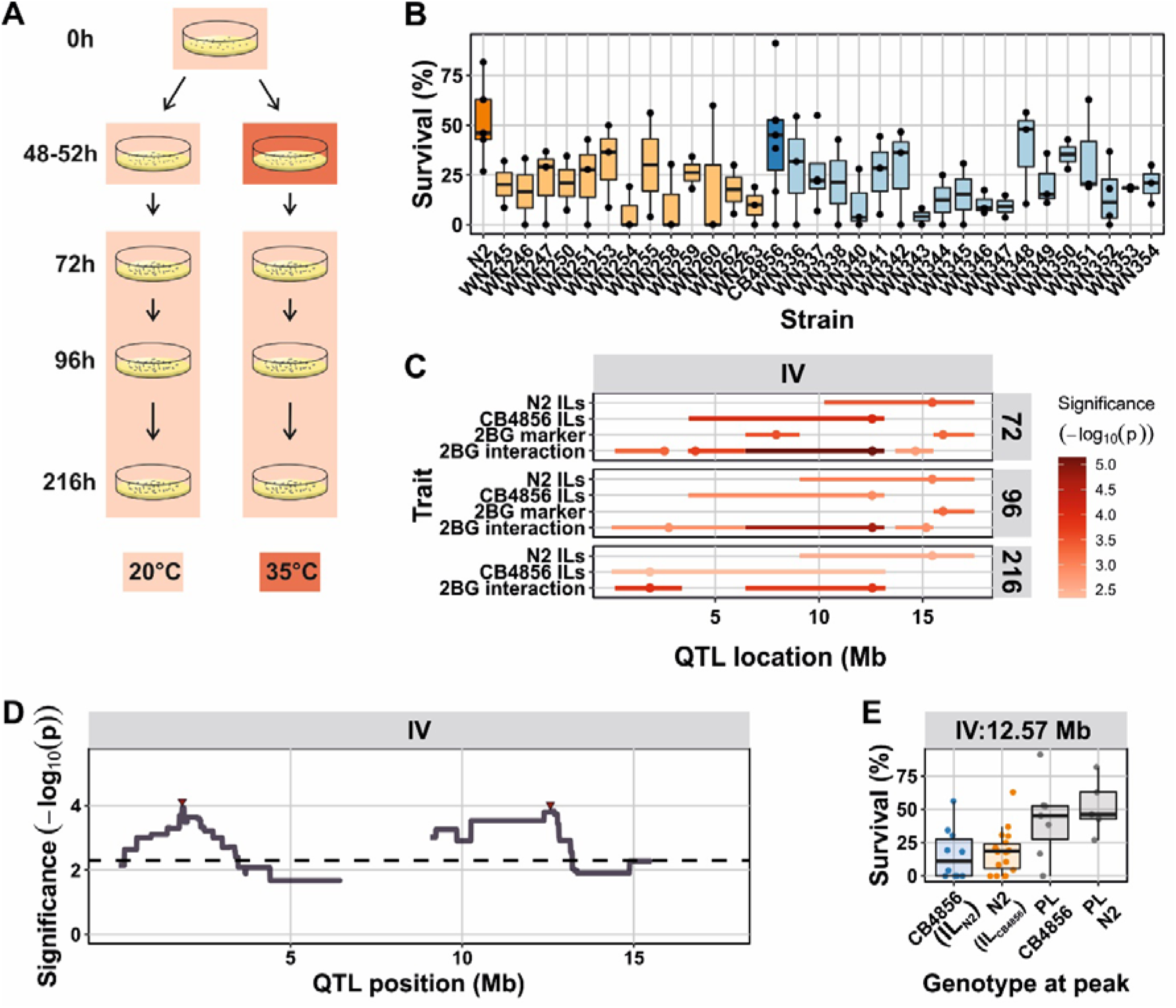
Heat-stress survival in ILs covering chromosome IV. (**A**) Experimental setup of the heat-stress survival experiment. (**B**) The survival one week after 4 hour exposure to a 35^°^C heat-shock. Each dot represents a plate (n = 20 – 40 animals per plate) and the box-plots are added as visual aide. The colours indicate the genetic background (orange: N2, blue CB4856). (**C**) Overview of all QTL mapped in the IL populations using various types of models (N2 ILs: IL_N2_ bin mapping; CB4856 ILs: IL_CB4856_ bin mapping; 2BG marker: 2-background mapping model variance captured by marker; 2BG interaction: 2-background mapping model variance captured by the interaction marker-background). On the x-axis, the physical position in million bases (Mb) is shown. The colour scale indicates the significance of the association. Only significant associations are shown (FDR = 0.05; based on 1,000 permutations). (**D**) The QTL profile of the 2BG interaction term, the two red triangles indicate the locations of the peaks. On the x-axis, the physical position in million bases (Mb) is shown, on the y-axis the significance. The dashed horizontal line indicates the threshold (FDR = 0.05; based on 1,000 permutations). (**E**) The QTL effects of the right peak from panel (D). Where both parental lines (PL) have a similar survival percentage, the IL_N2_ (CB4856 introgression at the locus) and the IL_CB4856_ (N2 introgression at the locus) display a lower survival percentage.

### Whole-genome QTL mapping uncovering an N2-background-dependent QTL for clec-62 gene-expression

One final way in which the IL_CB4856_ panel can be used, is by combining data from different genetic panels, especially the IL_N2_ and N2xCB4856 RIL populations. For this case, we investigated an expression QTL for the gene *clec-62*, which we previously picked up in the IL_N2_ panel (STERKEN *et al*. 2020). Previously, no expression QTL for *clec-62* in N2xCB4856 RILs had been identified (LI *et al*. 2010; ROCKMAN *et al*. 2010; SNOEK *et al*. 2017; STERKEN *et al*. 2017; SNOEK *et al*. 2020). We set out to verify previous QTL detection in the IL_N2_ panel (STERKEN *et al*. 2020), as well as explore the existence of eQTL for *clec-62* in the IL_CB4856_ population.

We measured the expression of the two isoforms of *clec-62* in 47 RILs, 42 IL_N2_ strains, and 41 IL_CB4856_ strains by RT-qPCR (**Figure 4A**; **Supplementary table 9**) and found that *clec-62A* expression was 3.5-fold higher than *clec-62B*. The expression levels of the two isoforms were highly correlated (**Supplementary figure 5A;** Pearson *R = 0.94*; p < 1^-15^) and therefore the analysis was performed using the higher expressed isoform A. We also found a high correlation between the expression in the previous microarray-based experiment on the IL_N2_ population, but not on the RIL population (**Supplementary figure 5B; Supplementary table 10**). This confirmed that we only found a QTL using the ILs. Analysis of broad-sense and narrow-sense heritabilities in the three populations showed there was only significant broad-sense heritability for *clec-62* expression in the IL_N2_ panel (*H^2^* estimates 0.86 – 0.98; q < 0.01; **Supplementary table 11**). Significant narrow-sense heritability was only found in the microarray data of the IL_N2_ panel (*h^2^* = 0.56; q < 0.01), which is probably due to the higher number of strains included in that experiment. Altogether, we hypothesize that genetic variation in *clec-62* expression can only be found in an N2 genetic background and no QTL should exist in the RIL and the IL_CB4856_ population. Therefore, the variation in *clec-62* expression must be explained by background interactions.

**Figure 4:**
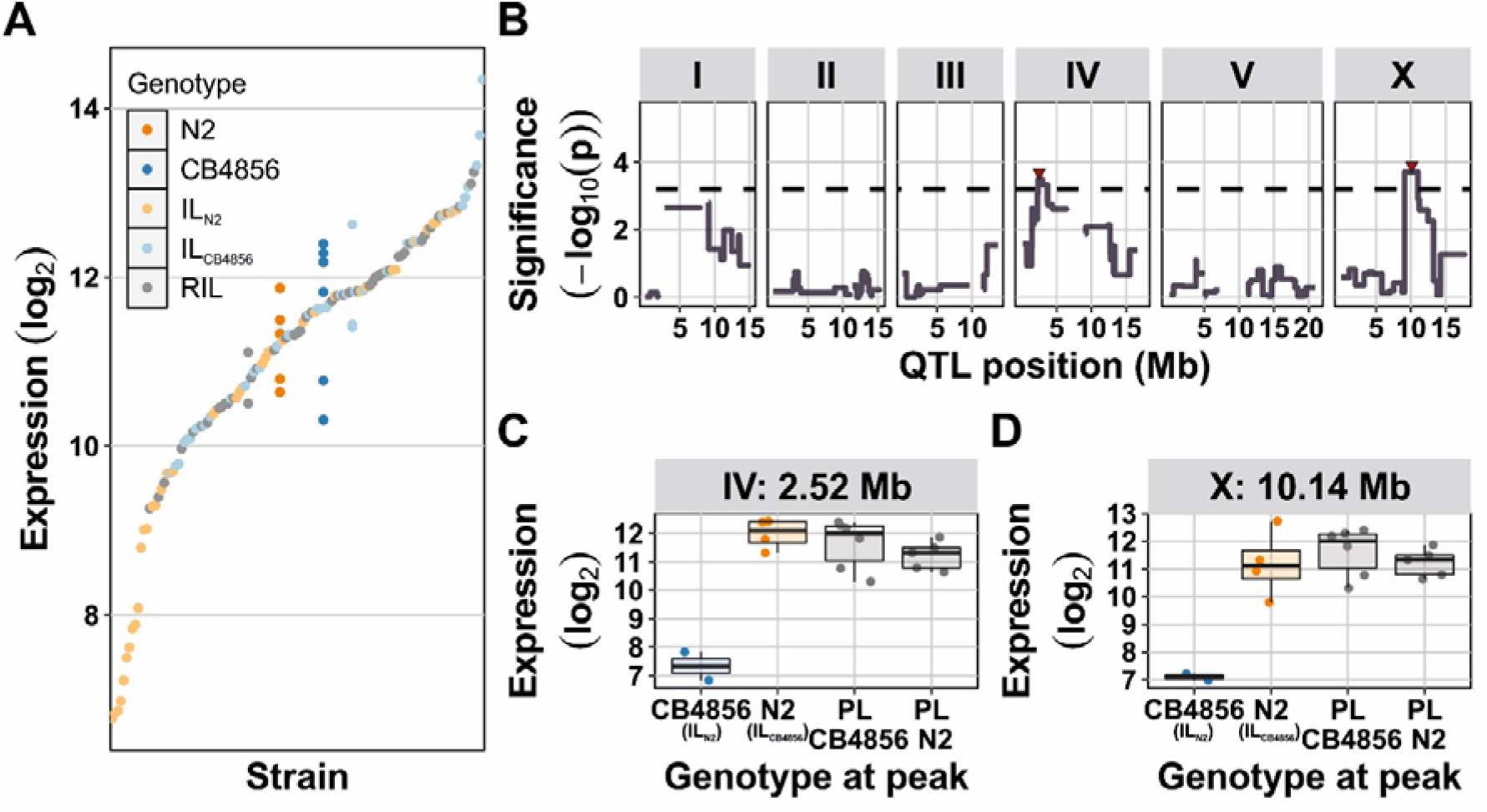
Expression of *clec-62* is due to QTL interacting with the N2 genetic background (**A**) The expression of *clec-62* as measured by qPCR in N2 (n = 5), CB4856 (n = 6), and three inbred panels: RIL (n =47), IL_N2_ (n = 42), and IL_CB4856_ (n = 41). The log_2_-normalized expression is shown per strain. (**B**) Profile of the significance of the interaction with the genetic background as mapped in the two IL panels. On the x-axis the physical position is shown in million bases (Mb). On the y-axis the significance of the association is shown (- log_10_(p)). The dashed horizontal line indicates the FDR = 0.05 threshold. The red triangles indicate QTL-peaks. (**C**) The split-out of the chromosome IV peak from panel (**B**). Where both parental lines (PL) and the IL_CB4856_ population have a similar *clec-62* expression, the IL_N2_ (CB4856 introgression at the locus) has a lower *clec-62* expression. (**D**) As in (**C**), but for the chromosome X peak.

QTL mapping of *clec-62* expression indeed identified QTL interacting with the genetic background. As predicted, no QTL were detected in the RIL and the IL_CB4856_ populations (**Supplementary figure 5C**). However, when mapping using the IL_N2_ panel, a highly similar QTL profile for *clec-62* expression as measured by RT-qPCR and microarray was found (**Supplementary figure 5C; Supplementary table 6**). When using the combined IL_N2_ and IL_CB4856_ panels, we identified two loci with significant interaction effects: one on chromosome IV and one on chromosome X (**Supplementary table 6; Figure 4B**). These loci were both associated with a low expression when a CB4856 introgression is present (**Figure 4C and D**). Overall, we conclude that the interaction of CB4856 introgressions with the N2 genetic background drives the *clec-62* expression QTL.

## Discussion

Here we presented a novel whole-genome N2>CB4856 IL population in *C. elegans*. This new IL_CB4856_ population is reciprocal to the previous IL_N2_ population (DOROSZUK *et al*. 2009). The construction of the novel panel was facilitated by the availability of the first high-quality CB4856 genome (THOMPSON *et al*. 2015), allowing for the selection of insertions/deletions as genetic markers which could be used during the crossing process. Genotyping by low-coverage whole-genome sequencing provided a detailed genetic map that allows the use of the novel population in various QTL approaches. We present three such approaches in this paper: (i) mapping lifespan using the panel on its own, (ii) further dissection of a previously implicated region in heat-stress resistance, and (iii) exploration of *clec-62* expression regulation using all three available N2xCB4856 populations. These cases show that the IL_CB4856_ panel empowers understanding the role of natural genetic variation in trait regulation, especially the role of genetic interactions.

### The IL_CB4856_ panel for use in QTL mapping

The IL_CB4856_ panel is mostly free from large-effect laboratory-derived alleles that segregate in the IL_N2_ panel. As the background of the developed IL population is CB4856, it lacks the N2 laboratory derived alleles, such as *nath-10* (DUVEAU AND FELIX 2012), *glb-5* (MCGRATH *et al*. 2009), and *npr-1* (MCGRATH *et al*. 2009; ANDERSEN *et al*. 2014). Therefore, the new IL_CB4856_ panel may be especially useful if the studied trait might be affected by these pleiotropic and large-effect alleles (STERKEN *et al*. 2015). Therefore, the crossing scheme used to obtain the IL population considered results obtained from quantitative genetics studies in *C. elegans*. The scheme accounted for the *peel-1*/*zeel-1* locus (SEIDEL *et al*. 2011) leading to marker distribution distortions in N2xCB4856 RIL populations (LI *et al*. 2006; ROCKMAN AND KRUGLYAK 2009; THOMPSON *et al*. 2015). A double back-cross with CB4856 was used to remove the N2 *peel-1/zeel-1* allele, which failed to segregate otherwise (data not shown).

One of the main strengths of IL populations in comparison to RIL populations lies in the detection of small phenotypic effects. RIL populations are hampered by the residual variance induced by the segregation of multiple QTL (KEURENTJES *et al*. 2007a). In strains with a homogeneous background all QTL are fixed, except the introgressed locus, reducing the residual variance (the experimental variation) per QTL to a minimum. Many studies observed ILs resolve more QTL than RILs (KEURENTJES *et al*. 2007a; GREEN *et al*. 2013; GLATER *et al*. 2014; BERNSTEIN AND ROCKMAN 2016; EVANS AND ANDERSEN 2020; EVANS *et al*. 2020) and because the IL_CB4856_ population contains novel breakpoints compared to the IL_N2_ population QTL in these ILs can now also be pinpointed to a narrower locus. Therefore, the IL_CB4856_ population will likely be useful for fine mapping complex traits. Furthermore, the IL_CB4856_ population can serve as a resource for the generation of ILs with even smaller introgressions (BERNSTEIN AND ROCKMAN 2016).

### The compounding benefit of additional mapping populations

We show that reciprocal IL populations can easily identify genetic interaction dependent on background-effects. As examples, we identified interacting loci for both heat-stress resistance and *clec-62* expression. The reason that genetic interactions can more easily be identified when compared to RIL population (BLOOM *et al*. 2015; FORSBERG *et al*. 2017) is that reciprocal IL populations can place the QTL effect in the context of the genetic background. There are many genome-wide IL populations, including for barley (SCHMALENBACH *et al*. 2008), *Arabidopsis* (KEURENTJES *et al*. 2007a; FLETCHER *et al*. 2013), tomato (MONFORTE AND TANKSLEY 2000), maize (SZALMA *et al*. 2007), rice (LI *et al*. 2005), and mice (GALE *et al*. 2009). However, to our knowledge there are no genome-wide reciprocal IL populations and until now observations in reciprocal ILs have only been made on a small scale. One difficulty of mapping interactions in reciprocal IL panels is that the partner locus (hiding in the genetic background) cannot readily be pinpointed. Still, QTL mapping in the combined IL_N2_ and IL_CB4856_ panels provides a starting point for investigations into genetic interactions.

From the experimental observations we made, we propose that genetic background effects are often overlooked in an IL panel with a single background. In such IL panels, the QTL-background interactions are confounded by definition. In other words: if an interaction with the genetic background exists and determines the trait levels in the individual introgression line, it cannot be distinguished from additive QTL. The homogenous background can lead to different estimations of QTL effect sizes compared to RILs (as reviewed (MACKAY 2014)). The cause of this effect is due to the frequency of the genotype at the interacting locus. In an ideal RIL population, the loci are unlinked and therefore both genotypes affect the main effect at the QTL. However, in ILs, the loci are linked and the QTL main effects are therefore affected by the background interaction as exemplified by ILs generated for specific loci (KROYMANN AND MITCHELL-OLDS 2005; MCGRATH *et al*. 2009; GAERTNER *et al*. 2012). The availability of two reciprocal IL populations therefore enables QTL dissection on a genome-wide scale.

## Acknowledgements

The authors thank Robyn Tanny, Erik Andersen, Beatrice Tan, Tatiana Blokhina, Jenifer Sundar, and Myrthe Walhout for technical support. Erik Andersen is also thanked for feedback on the manuscript. MGS was supported by the Graduate School for Production Ecology and Resource Conservation (PE&RC) and by NWO domain Applied and Engineering Sciences VENI grant (17282). LBS was funded by ALW grant 823.01.001 JEK received support from NIH R01 AA026658.

## Author contributions

Conceived and designed the experiments: MGS, LBS, JEK. Constructed the panel: MGS, JWvC, JAGR. Sequencing and genetic map construction: DC. Performed experiments: MS, JS, KJ, YAW, JJS, SCH. Analysed the data: MGS, LvS. Wrote the paper: MGS, LvS. All authors edited and commented on the paper.

## Supplementary figures and tables

**Supplementary figure 1**: Map of the amplification fragment length polymorphisms. For each primer-pair (indicated by the chromosome number and the start location of the deletion in CB4856) the amplicons in N2, CB4856 and the six RILs used for constructing the IL_CB4856_ panel are shown. The 100, 500, and 1000 bp bands of the 1kb+ marker are indicated. Photographs of the gels were stretched to cover the same area.

**Supplementary figure 2**: Power analysis in the IL_CB4856_ and the combined IL_N2_ and IL_CB4856_ panels. (**A**) Power analysis of bin mapping using the IL_CB4856_ panel core set of 87 ILs. The percentage of simulated QTL detected is plotted versus the number of replicates per IL. The colours indicate the amount of heritable variation simulated in the QTL. (**B**) Power analysis of bin mapping using both the 90 IL_N2_ and 87 IL_CB4856_ core sets. The simulated QTL contained only a marker effect. The percentage detected versus the number of replicates is for the marker-effect in the two-background model.

**Supplementary figure 3**: A figure of all lifespan QTL mapped. On the y-axis the various summary statistics (maximum, mean, median, and variance) for lifespan are shown. On the x-axis the position of the QTL in million bases (Mb) is shown. The dots indicate the peak of the QTL and the horizontal lines the confidence interval of the peak location. The colours indicate the significance.

**Supplementary figure 4:** The heat-stress survival QTL mapped in the data obtained from (JOVIC *et al*. 2019). (**A**) the QTL profiles as mapped in the IL_N2_ and the N2xCB4856 RIL population. On the x-axis the physical position is shown in million bases (Mb). On the y-axis the significance of the association is shown (-log_10_(p)). The dashed horizontal line indicates the FDR = 0.05 threshold. The red triangles indicate QTL-peaks. (**B**) The split-out per genotype at the Chromosome IV QTL at 13.2 Mb. The ILs with the CB4856 introgression at the marker are shown versus the measurements in the N2 strain.

**Supplementary figure 5:** The *clec-62* expression QTL. (**A**) The expression of *clec-62* isoform A versus isoform B as measured by RT-qPCR in the same strains. The expression has been normalized and log_2_-transformed. The Pearson correlation coefficient is shown as well as the significance of correlation. The colours indicate the various strains/panels tested. (**B**) the correlation between the expression of *clec-62* as measured by microarray (x-axis) versus RT-qPCR measured *clec-62* isoform A expression. Each dot represents the expression measured in a strain of either the IL_N2_ panel (orange) or RIL panel (grey). The Pearson correlation coefficient is shown, as well as the significance of the correlation. (**C**) The QTL profiles of *clec-62* expression. On the x-axis the physical position is shown in million bases (Mb). On the y-axis the significance of the association is shown (-log_10_(p)). The dashed horizontal line indicates the FDR = 0.05 threshold. The red triangles indicate QTL-peaks. Where qPCR data is shown, *clec-62* isoform A was used for analysis.

**Supplementary table 1:** Description of the crossing setup and generations underlying each IL.

**Supplementary table 2:** List with the fragment length polymorphism primers used for initial tracking of segments.

**Supplementary table 3:** Complete list of genotypes of all the Wageningen Nematology strains of the N2xCB4856 RIL, IL_N2_ and IL_CB4856_ panels.

**Supplementary table 4:** Integrated genetic map for all the Wageningen Nematology strains of the N2xCB4856 RIL, IL_N2_ and IL_CB4856_ panels.

**Supplementary table 5:** Data of the lifespan experiment in the IL_CB4856_ panel.

**Supplementary table 6:** Locations and intervals of all QTL mapped for lifespan, heat-stress survival and *clec-62* expression.

**Supplementary table 7:** Heat stress survival data from (JOVIC *et al*. 2019) that was analysed.

**Supplementary table 8:** Heat stress survival data from chromosome IV IL_N2_ and IL_CB4856_ strains.

**Supplementary table 9:** Expression data of the *clec-62* gene as measured by RT-qPCR.

**Supplementary table 10:** Expression data of the *clec-62* gene as measured by microarray, obtained from (SNOEK *et al*. 2017; STERKEN *et al*. 2020).

**Supplementary table 11:** Heritability analysis of *clec-62* expression.

## Notes

https://www.ncbi.nlm.nih.gov/sra/?term=SRP154243

### Competing Interest Statement

The authors have declared no competing interest.

### Summary of Updates

Clarification of axes of main figures (2, 3, 4) and supplementary figure 4. Additionally, adaptions were made in the text to further clarify some parts.

