## Supplementary figures and images for "Two sides to every coin: reciprocal introgression line populations in *Caenorhabditis elegans*"

### Supplementary figure 1

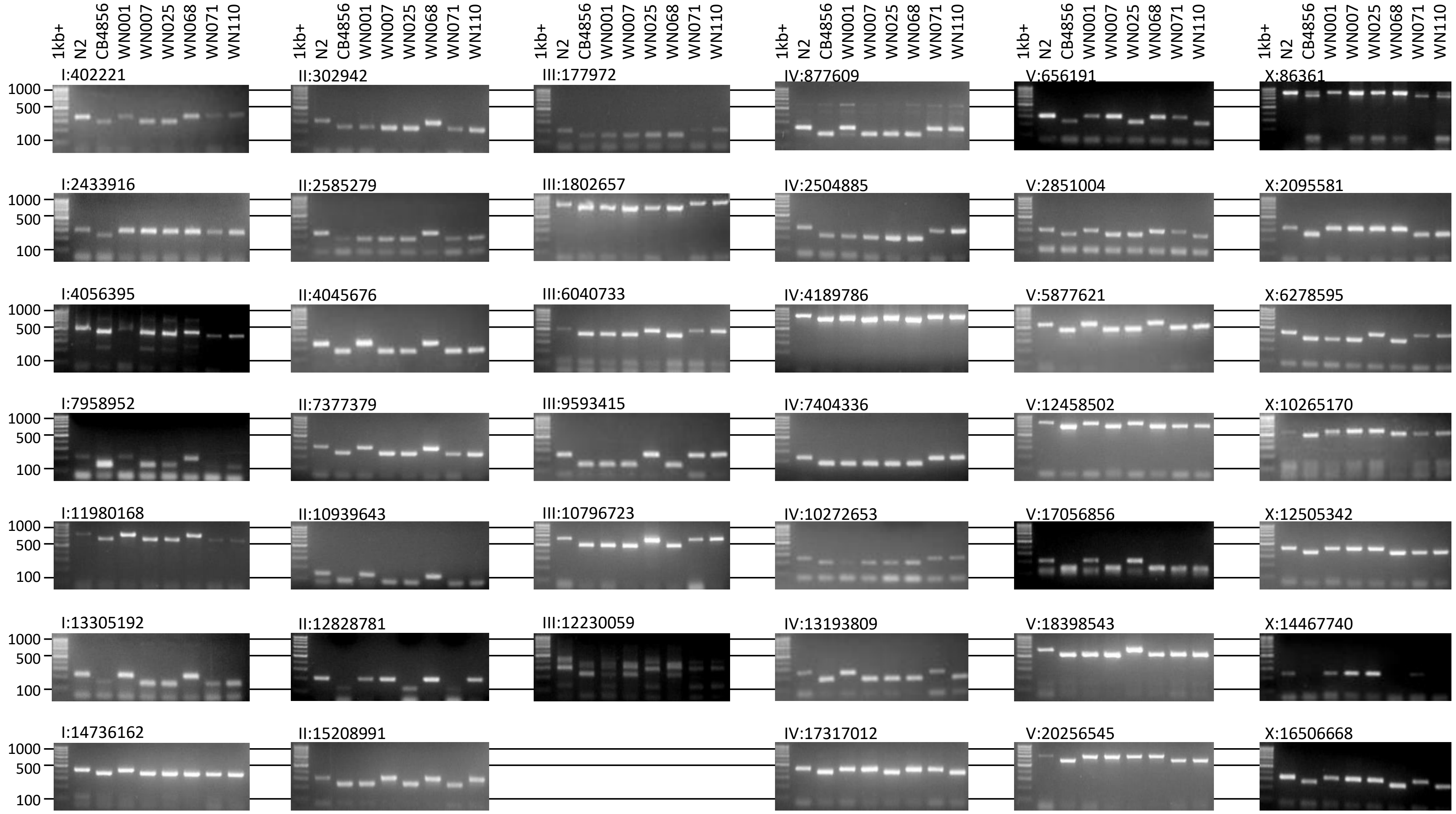

### Supplementary figure 2

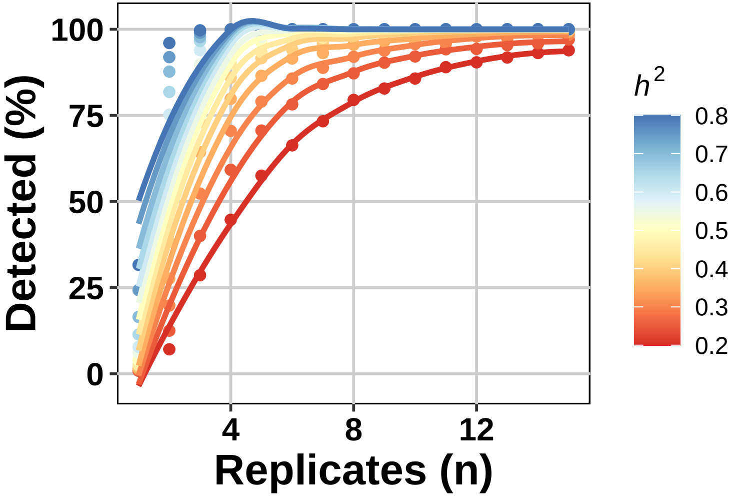

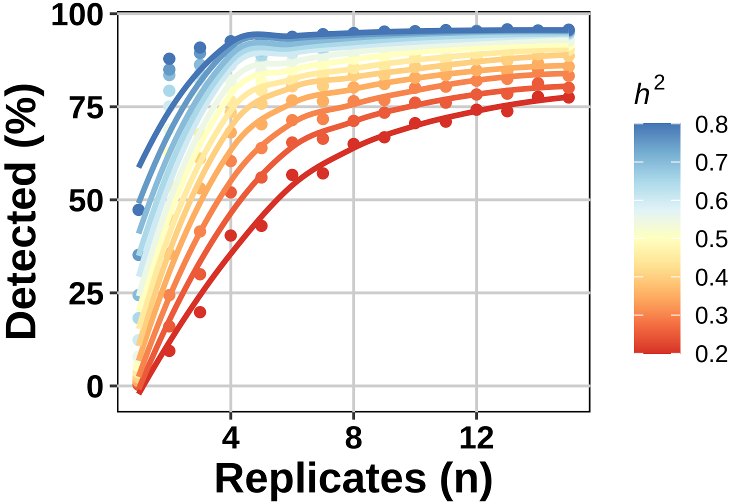

### Supplementary figure 3

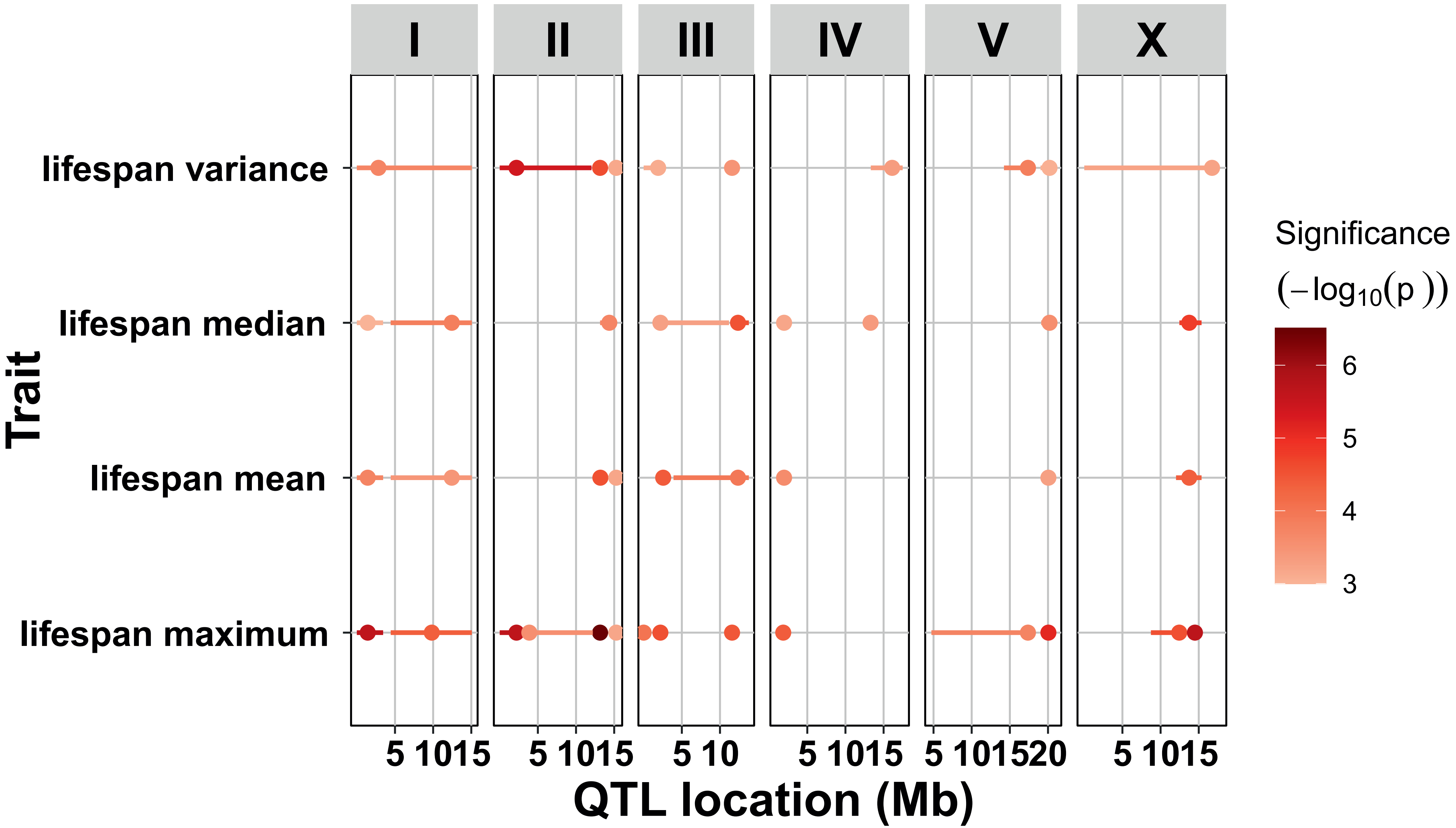

### Supplementary figure 4

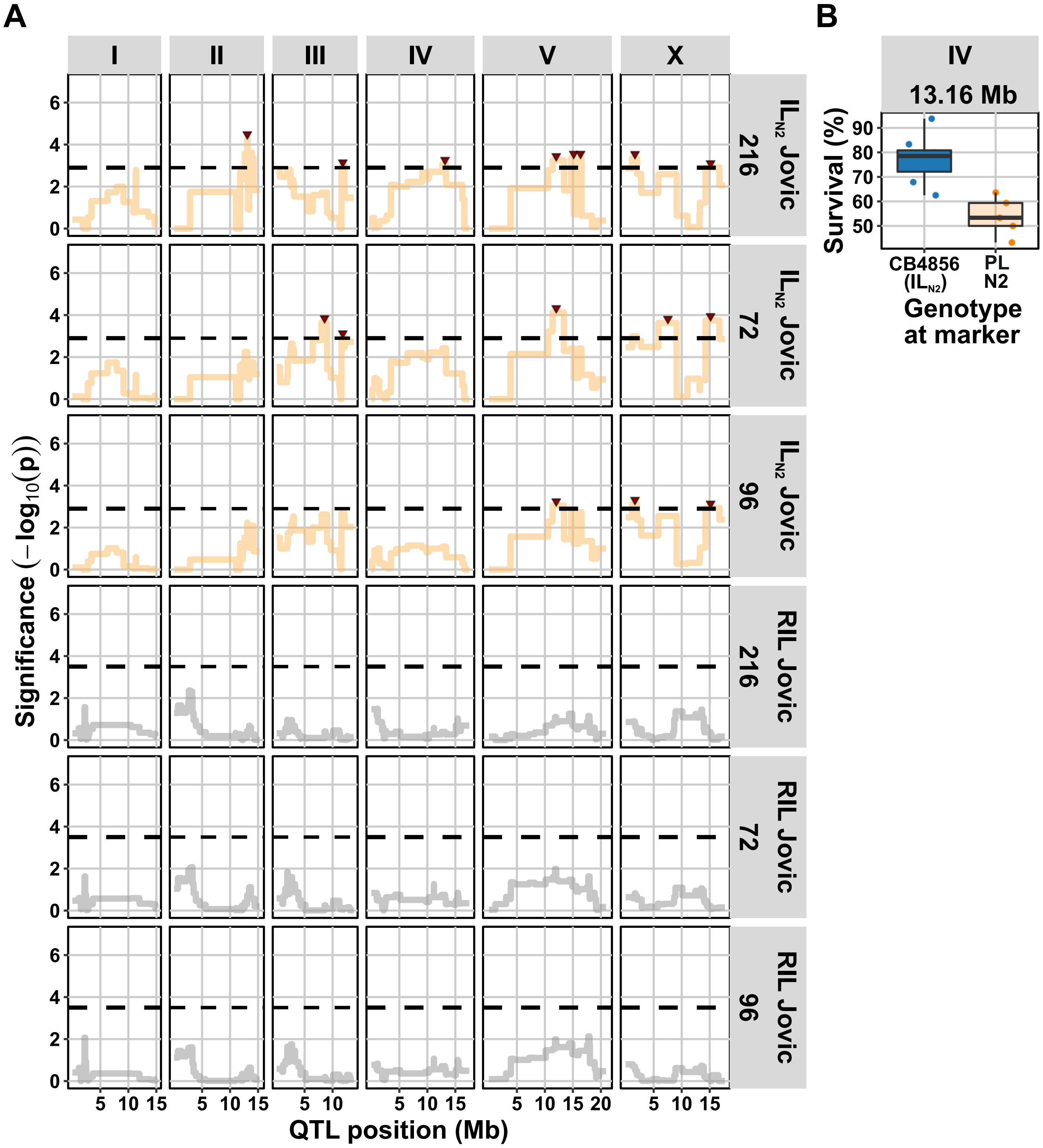

### Supplementary figure 5

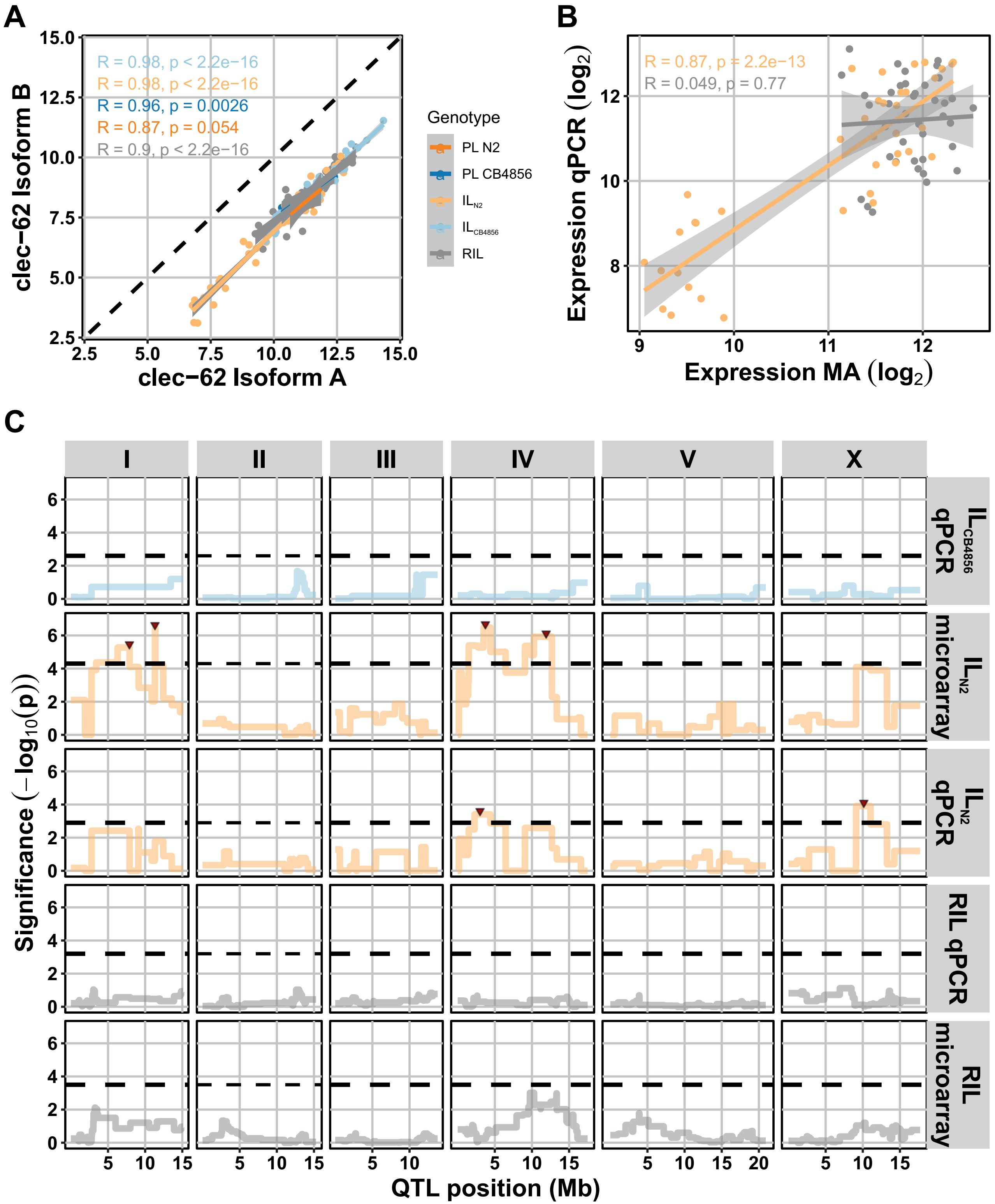
